# Context Shapes Fear: Habituation of Innate Defensive Behaviours Depends on Environmental Context

**DOI:** 10.1101/2025.08.21.671638

**Authors:** Haoxuan Qi, Alice Treloar, Greg J. Stuart, Saba Gharaei

## Abstract

The capacity of animals to rapidly and appropriately respond to potential threats is critical for survival. In many species, this involves innate defensive behaviours, such as escape or freezing. But not all threats are dangerous. Habituation – a form of non-associative learning – allows animals to filter out irrelevant stimuli and thereby avoid unnecessary energy expenditure. While it is well known that context can influence behavioural responses to associative learning, whether non-associative forms of learning, such as habituation, are also context-dependent is unclear. Here, we address this issue in mice by examining the role of environmental context on habituation of defensive behaviours to threatening visual stimuli. We first developed a protocol leading to rapid (within minutes) and stable (at least one week) habituation of freezing responses to slowly sweeping visual stimuli, resembling an aerial predator moving across the sky. Using this protocol, we tested the impact of environmental context on habituation of freezing responses, finding that changing the environmental context led to a significant and reversible reduction in habituation. In summary, we find that environmental context plays a critical role in determining the impact of habituation on visually evoked innate defensive behaviours. These findings reveal a previously unrecognised flexibility in defensive responses to visual threats, extending the influence of environmental context to non-associative learning.

**Highlights:**

- Mice freeze in response to sweeping visual stimuli, resembling an aerial predator.
- Habituation to sweeping visual stimuli could be induced rapidly and remained stable over time.
- Habituation to sweeping visual stimuli depended on environmental context.
- Context-specific habituation was reversible.

## Introduction

Survival in the natural world requires animals to rapidly and appropriately respond to potential threats^1–3^. Across multiple species, this has led to the evolution of innate defensive behaviours that can be automatically triggered by biologically relevant threatening stimuli^1,4–8^. These behaviours are distinct from conditioned fear responses in that they do not depend on prior learning, suggesting that they are mediated by hard-wired neural circuits designed to detect and react to threats with minimum processing delay^9^.

The type of defensive behaviour an animal expresses depends on the nature of the perceived threat. According to the threat imminence theory, specific sensory cues are matched to specific defensive strategies^7^. For example, slowly sweeping visual stimuli, resembling an aerial predator moving across the sky, typically elicit freezing in rodents, presumably to reduce detection^10,11^. In contrast, looming visual stimuli, which simulate a rapidly approaching aerial predator, usually trigger escape^10–13^. While these responses appear stereotyped, emerging evidence suggests that innate defensive behaviours are not fixed and can exhibit a degree of plasticity depending on external conditions and prior experience^14–17^.

One important form of plasticity of innate behaviours is habituation, where repeated exposure to a non-threatening stimulus leads to a progressive reduction in the behavioural response^15,17–19^. Habituation of innate responses to sensory input enables organisms to filter out irrelevant stimuli and thereby avoid unnecessary energy expenditure^20^. With regard to habituation to visual threats, repeated exposure to looming stimuli can rapidly reduce escape behaviour in mice within minutes^15,17,18,21^. Exposing mice to multiple sweeping stimuli has also been shown to reduce freezing responses^17,19^. While these studies establish that mice can undergo habituation to visual threats, it is not known how the context in which a visual threat is presented influences habituation of defensive behaviours to threatening visual stimuli.

The role of environmental context in associative forms of learning, such as fear conditioning, is well established^22–27^. In contextual fear conditioning, animals associate a specific environment with an aversive event (e.g. an electric shock), triggering freezing. Re-exposure to the same environment triggers freezing, whereas exposure to a novel environment does not^23–25^. These findings demonstrate that environmental cues play a crucial role in modulating defensive responses in this associative form of learning. Whether non-associative forms of learning, such as habituation of innate defensive behaviours triggered by visual threats, are also dependent on context is unknown.

In this study, we developed a novel behavioural paradigm that rapidly (within minutes) induces habituation of freezing responses to sweeping visual stimuli in mice. Habituation induced by this paradigm was stable over time (up to at least one week). Using this habituation paradigm, we systematically examined the factors that influence the expression and stability of habituation to threatening visual stimuli. We found that environmental context plays a critical role in determining the impact of habituation on visually evoked defensive behaviours. These experiments provide new insight into how environmental context shapes habituation of innate defensive behaviours and its role in non-associative learning.

## Results

### Rapid Habituation to Visual Sweeping Stimulus

We first developed a behavioural habituation protocol to rapidly habituate mice to sweeping visual stimuli. This protocol was based on one recently developed to generate habituation to looming visual stimuli^18^. Mice were placed in an arena with a monitor mounted overhead^28^. The monitor was used to display a small black disk sweeping overhead to simulate a cruising aerial predator (Figure 1A). Mice were randomly assigned to a control or habituated group and only tested with the sweeping stimulus if/when they entered the centre of the arena (marked by four dots on the floor). Mice in the control group were allowed to explore the arena for 9 minutes with no overhead visual stimulus. In contrast, mice in the habituated group were exposed to 130 repetitions of the visual sweeping stimulus over the same 9-minute period, during which the screen’s background gradually brightened, slowly enhancing the contrast between the sweeping stimulus and the background. Following this 9-minute period, we tested the behavioural response of both groups to three full-contrast sweeping stimuli (Figure 1B).

**Figure 1.**
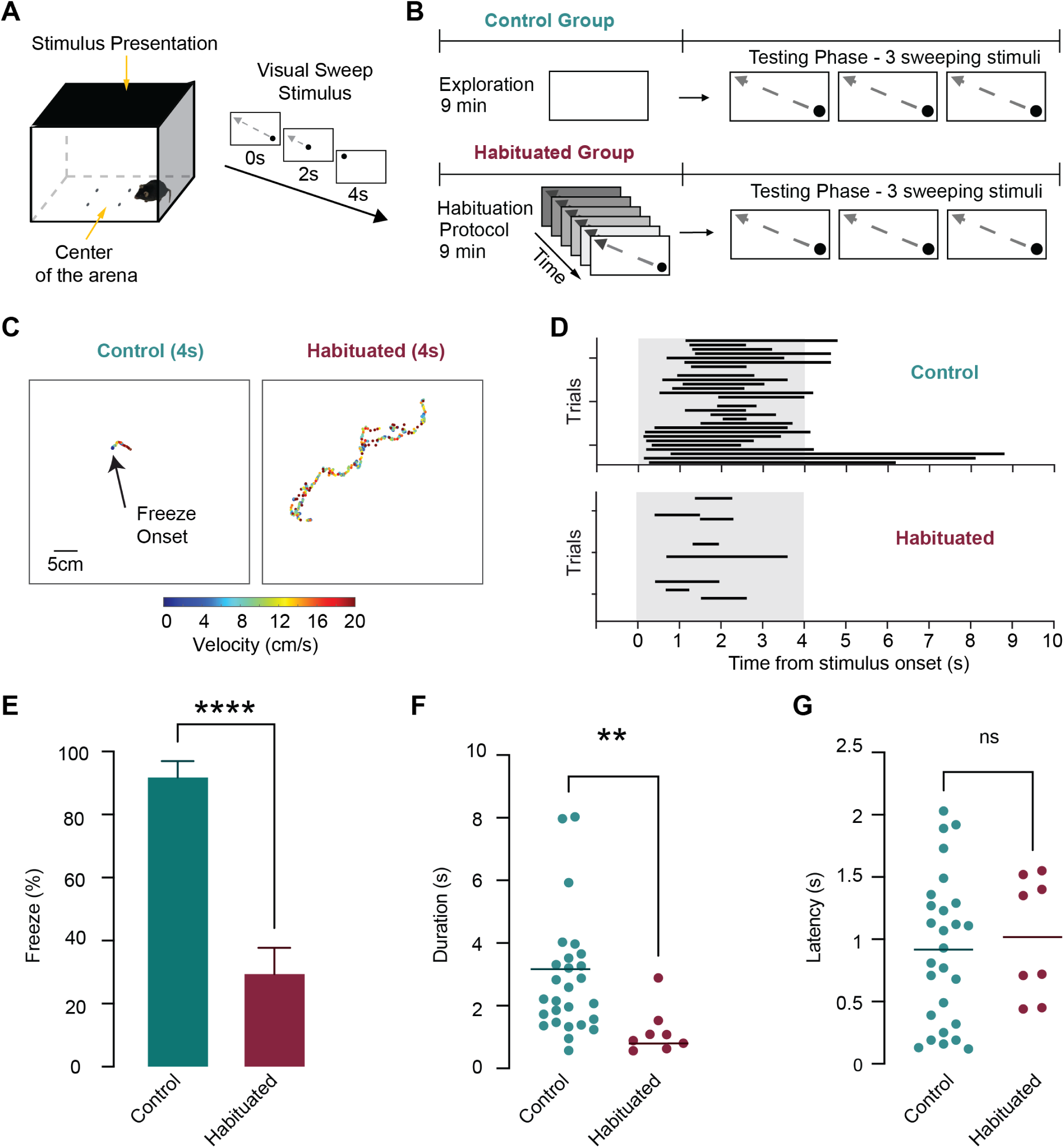
Repeated exposure to sweeping visual stimuli rapidly suppresses innate freezing responses. (A) Left, schematic of the testing arena used for the habituation protocol. Right, the visual sweeping stimulus is a small black disk that sweeps diagonally across the overhead monitor over 4 seconds. (B) Experimental design. Mice were randomly assigned to the control group (9-minute exploration with no visual stimuli) or the habituation group (130 sweeping visual stimuli increasing in contrast over 9 minutes). Both groups were subsequently tested with three identical full-contrast sweeping stimuli. (C) Representative DeepLabCut trajectory plots showing mouse movement speed and position in the control and habituated groups during the 4 seconds of the test sweep stimulus. (D) Raster plots showing trial-by-trial freezing responses to the test stimuli in control (teal) and habituated mice (burgundy). Black bars indicate freezing. The shaded grey area indicates stimulus presentation. (E) Average percentage of trials (±SEM) where freezing was observed (across mice) in the control (teal) and habituated group (burgundy). Habituated mice froze significantly less than controls. (F, G) Freeze duration (F) and latency (G) in control (teal) and habituated mice (burgundy). Horizontal lines indicate the median. Habituated mice showed significantly shorter duration freezing episodes, but with similar onset latencies.

The vast majority of mice in the control group froze in response to the test sweep stimulus, whereas freezing was largely absent in the habituated group. In the control group, freezing was observed in 27 out of 29 trials, compared to just 8 out of 31 trials in the habituated group (Figure 1C–D; 11 mice per group; up to 3 trials per mouse). On average, the percentage of trials resulting in freezing was significantly reduced in the habituated group compared to control animals (Figure 1E; Control: 92.5 ± 5.2%; Habituated: 30.1 ± 8.1%; unpaired t-test, p < 0.0001; n=11 mice per group). Habituated mice also froze for significantly shorter durations on the trials where they did freeze (Figure 1F; Control median: 2.21s, IQR=1.47-5.32 s, n=11 mice; Habituated median: 0.99s, IQR=0.67-1.42 s, n=8 mice; Mann–Whitney test, p=0.0015). The time to freezing onset (freeze latency) did not differ significantly between the two groups (Figure 1G; Control: 0.92s ± 0.11s, n=11 mice; Habituated: 1.02s ± 0.17s, n=8 mice; unpaired t-test, p=0.6677). Together, these results demonstrate that gradual exposure to a sweeping visual stimulus can rapidly (within minutes) induce habituation of innate freezing responses in mice, reflected by reduced frequency and duration of freezing behaviour.

### Habituation Memory Lasts for at Least One Week

If habituation to non-threatening visual stimuli is to be behaviourally relevant, it should remain stable over time. To assess the stability of habituation, mice were tested at three time points: on the same day as the habituation protocol (Day 0), 24 hours later (Day 1) and one week later (Day 7). The percentage of trials resulting in freezing remained low across all three time points, with no significant difference in the extent of freezing across the days tested (Figure 2A; Day 0: 33.1 ± 12.7%, n=7 mice; Day 1: 41.5 ± 15.1%, n=8 mice; Day 7: 38.0 ± 8.8%, n=7 mice; one-way ANOVA, F(2, 19)=0.1126, p=0.8941). Similarly, freeze duration did not differ significantly between any of the test days (Figure 2B; Day 0 median: 0.95 s, IQR=0.72-1.10 s, n=5 mice; Day 1 median: 0.79 s, IQR=0.63-2.17 s, n=5 mice; Day 7 median: 1.29 s, IQR=0.71-1.62 s, n=6 mice; Kruskal-Wallis test, H=0.3551, p=0.8373, n=16 mice). Freeze latency also remained stable over time, with no significant differences observed between test days (Figure 2C: Day 0 median: 0.96 s, IQR=0.62-1.39 s, n=5 mice; Day 1 median: 0.99 s, IQR=0.94-2.35 s, n=5 mice; Day 7: 1.01 s, IQR=0.69-2.52 s, n=6 mice; Kruskal–Wallis test, H = 1.557, p = 0.4591, n = 16 mice). Together, these results indicate that habituation to sweeping visual stimuli is robust and persists for at least one week following a single habituation session.

**Figure 2.**
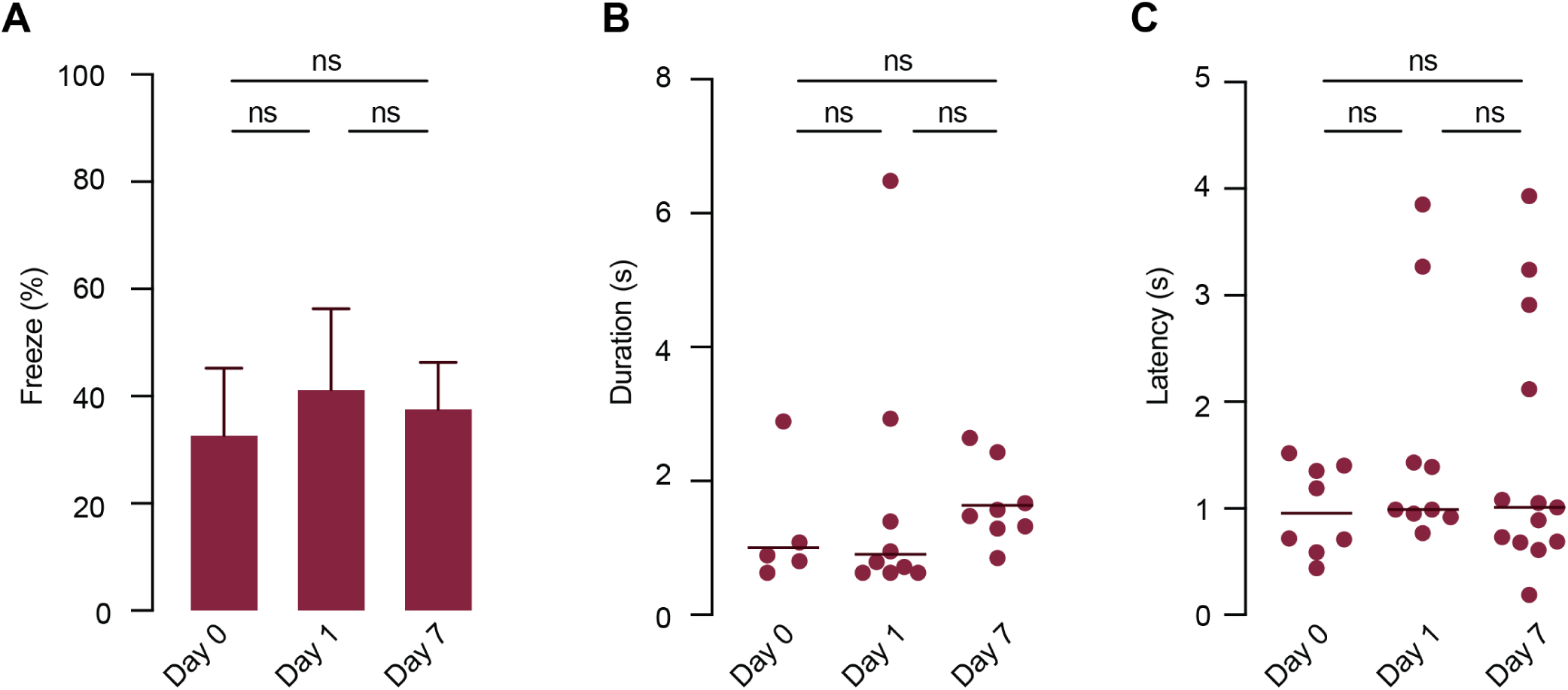
Habituation to sweeping visual stimuli is stable for at least one week. (A) Average percentage of trials (±SEM) where freezing was observed (across mice) on different days after the habituation protocol. Immediately after the habituation protocol (Day 0), 24 hours later (Day 1) and one week later (Day 7). No significant differences were observed across days. (B, C) Freeze duration (B) and latency (C) on the different testing days. Horizontal lines indicate the median. No significant differences were found across days.

### Habituation to Sweeping Stimuli Generalises Across Stimulus Direction

To test whether habituation was specific to the direction of the habituating sweeping stimulus, mice were habituated to a sweeping stimulus in one direction and then tested using a sweeping stimulus that was either in the same or the orthogonal direction (Figure 3A). The percentage of trials resulting in freezing did not differ significantly between these two test groups (Figure 3B; Same direction: 34.5 ± 8.3%, n=13 mice; Orthogonal direction: 30.7 ± 8.8%, n=13 mice; unpaired t-test, p=0.7542). Similarly, no significant difference was observed in freeze duration (Figure 3C: Same direction median: 0.91 s, IQR=0.62-1.23 s, n=9 mice; Orthogonal direction median: 0.97 s, IQR=0.6-1.27 s, n=8 mice; Mann–Whitney test, p=0.8918) or in freeze latency (Figure 3D; Same direction median: 1.02 s, IQR=0.71-1.65 s, n=9 mice; Orthogonal direction median: 2.32 s, IQR=0.71-3.01 s, n=8 mice; Mann–Whitney test, p=0.1693). These results indicate that habituation to sweeping stimuli does not depend on the direction of the sweeping stimulus.

**Figure 3.**
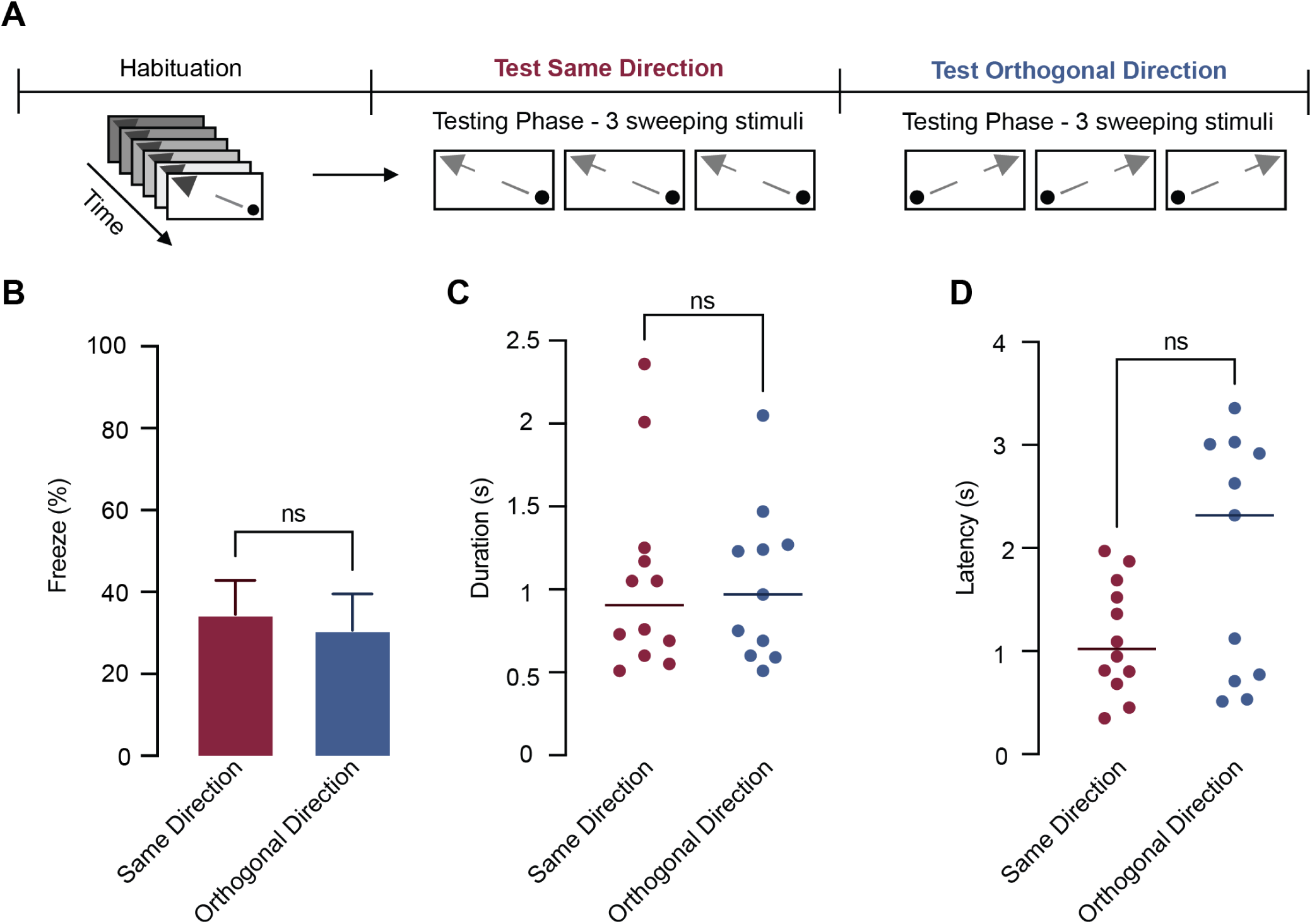
Habituation does not depend on the direction of the stimulus. (A) Schematic of the experimental design. After habituation, mice were tested with three sweeping stimuli either in the same direction as during habituation or in an orthogonal direction. (B) Average percentage of trials (±SEM) where freezing was observed (across mice) when tested with a sweeping stimulus presented in the same (burgundy) or the orthogonal (blue) direction to that used during habituation. Horizontal lines indicate the median. No significant differences were observed between groups. (C, D) Freeze duration (C) and latency (D) when mice were tested with a sweeping stimulus presented in the same (burgundy) or the orthogonal direction (blue) to that used during habituation. Horizontal lines indicate the median. Freezing duration and latency did not depend on stimulus direction.

### Habituation to Sweeping Stimuli is Context-Specific

We next tested whether habituation of innate defensive responses to sweeping visual stimuli depends on the environmental context. To test this, mice were habituated in one of two distinct contexts: Context A, which featured unique visual cues on the arena wall, and Context B, which had no visual cues on the arena walls and bubble wrap on the floor to provide tactile input (Figure 4A). Following habituation, mice were tested in either the same context in which they were habituated or in a different context (Figure 4B). The visual sweeping stimuli used for testing were identical in both groups. Mice tested in the same context in which they were habituated showed markedly less freezing compared to mice tested in a different context (Figure 4C; Same context: 29.6 ± 8.7%, n=9 mice; Different context: 89.0 ± 5.5%, n=9 mice; unpaired t-test, p < 0.0001). Despite this difference in freezing, there was no difference in freeze duration (Figure 4D, Same context median: 1.28 s, IQR=1.06-1.63 s, n=6 mice; Different context median: 1.14 s, IQR=0.77-1.54 s, n=9 mice; Mann–Whitney test, p=0.375) or latency between mice tested in the same or the different context from that used during habituation (Figure 4E, Same context median: 0.64 s, IQR=0.38-2.12 s, n=6 mice; Different context median: 0.35 s, IQR=0.13-0.82 s, n=9 mice; Mann–Whitney test, p=0.1507).

**Figure 4.**
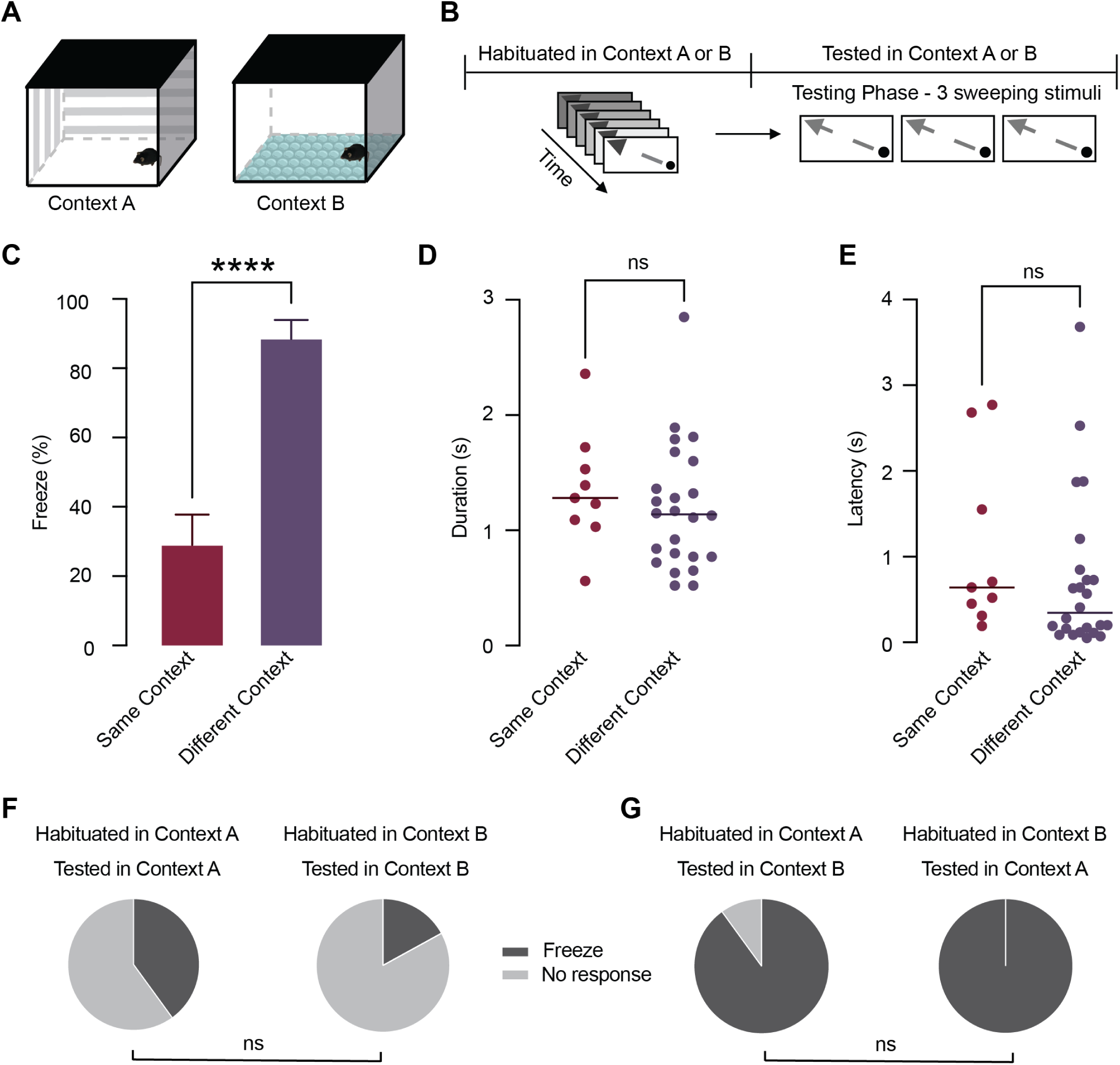
Habituation is context specific. (A) Illustration of the two environmental contexts used. Context A featured visual landmarks on the arena walls, while Context B had no visual landmarks and a bubble wrap floor. (B) Schematic of the experimental design. Mice were habituated in either Context A or B and then tested with three sweeping stimuli in either the same or a different context. (C) Average percentage of trials (±SEM) where freezing was observed (across mice) when tested in the same (burgundy) or a different context (purple) to that used during habituation. Freezing was significantly lower when mice were tested in the same context. (D, E) Freeze duration (D) and latency (E) when mice were tested in the same (burgundy) or in a different context (purple) to that used during habituation. Horizontal lines indicate the median. There was a significant difference between mice tested in the same or a different context. (F) Pie charts showing there was no significant difference in the proportion of freezing trials in mice habituated in Context A and tested in Context A, compared to mice habituated in Context B and tested in Context B. (G) Pie charts showing there was no significant difference in the proportion of freezing trials in mice habituated in Context A and tested in Context B, compared to mice habituated in Context B and tested in Context A.

To ensure that this context-specific effect was not due to intrinsic differences in threat salience between the two contexts used, we separated the data based on which context was used for habituation. There was no significant difference in the percentage of trials resulting in freezing when mice were habituated in Context A compared to Context B (Figure 4F; Context A: 40%, n=7 mice; Context B: 17% freezing, n=2 mice; Fisher’s exact test, p=0.3798). Similarly, the percentage of trials resulting in freezing when mice were tested in a context different from that used during habituation did not depend on whether mice were habituated in Context A or Context B (Figure 4G; Context A to B: 90% freezing, n=7 mice; Context B to A: 100% freezing, n=2 mice; Fisher’s exact test, p > 0.9999). Together, these results indicate that habituation to sweeping visual stimuli is context-specific, indicating that contextual cues play a crucial role in gating the habituated state.

### Context-Specific Habituation is Reversible

While these data show that habituation depends on context, it is unclear whether this effect reflects a general suppression of freezing in the new context or if there is an associative link that is specific to the habituation context. To address this, we performed a context reversal experiment to determine whether the reduction in habituation in the new context could be reversed by subsequent exposure to the same context in which mice were habituated (Figure 5A). Mice were first habituated in either Context A or Context B. They were then tested in either the same or a different context (Day 0). The following day (Day 1), the test context was reversed relative to that used on Day 0. On Day 5-9 mice were tested in the same context used immediately after habituation on Day 0. This design allowed us to determine whether returning to the initial habituation context restored the expression of habituation. That is, whether habituation is both context-specific and reversible.

**Figure 5.**
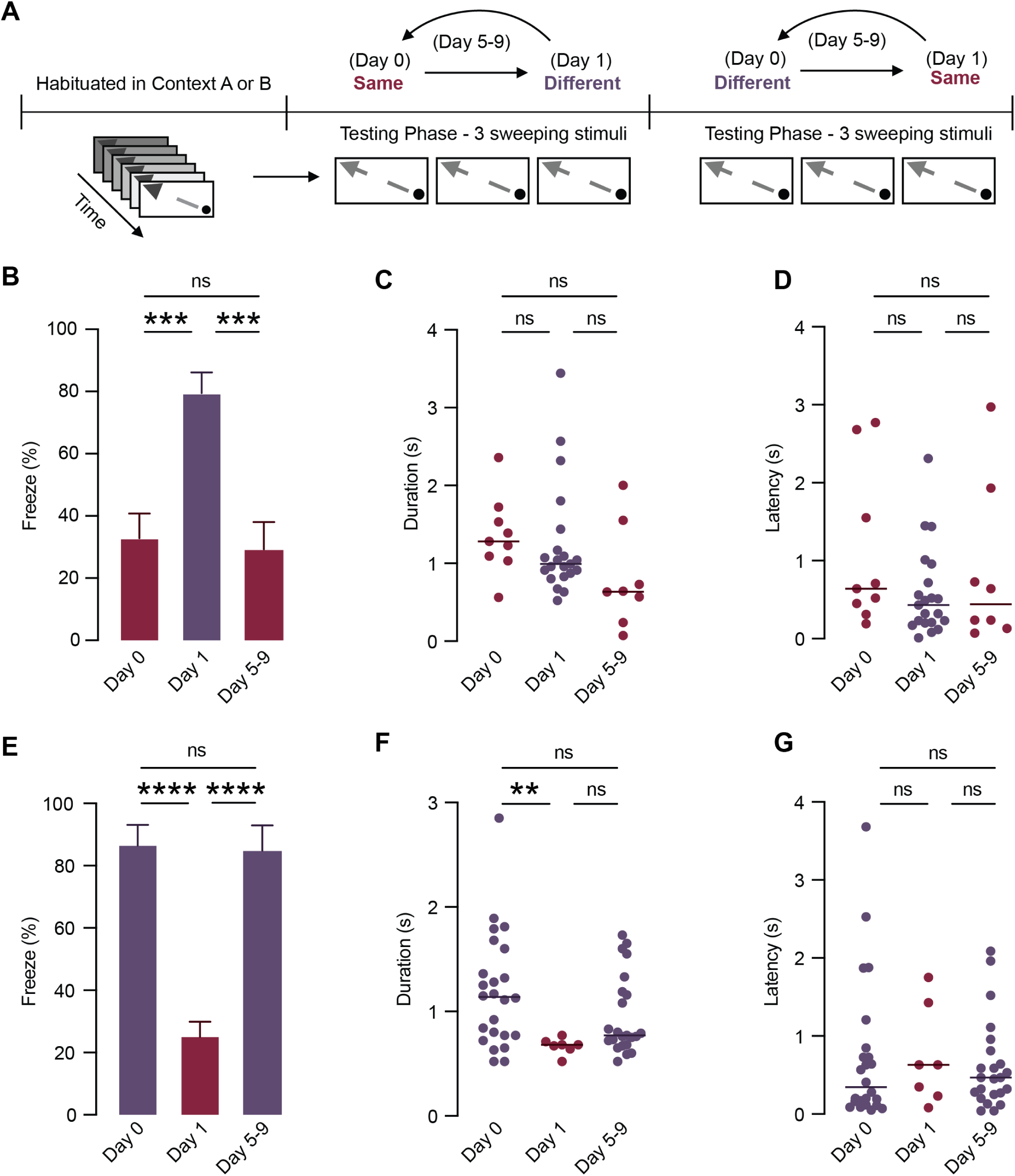
Context reversal reveals that habituation expression is gated by environmental context. (A) Experimental design for the context reversal paradigm. Mice were habituated in either Context A or B and tested across three time points: Day 0 (same or different context), Day 1 (context reversed from Day 1), and Day 5-9 (returned to Day 0 context). (B) Average percentage of trials (±SEM) where freezing was observed (across mice) on different days. Mice were initially tested in the same context as used during habituation (Day 0, burgundy), then tested in a different context on Day 1 (purple) and returned to the original Day 0 context on Day 5-9 (burgundy). (C, D) Freeze duration (C) and latency (D) in mice initially tested in the same context as used during habituation (Day 0, burgundy), then tested in a different context on Day 1 (purple) and then returned to the original Day 0 context on Day 5-9 (burgundy). Horizontal lines indicate the median. There was no significant difference between testing days. (E) Average percentage of trials (±SEM) where freezing was observed (across mice) on different days. Mice were initially tested in a different context to that used during habituation (Day 0, burgundy), then tested in the habituated (same) context on Day 1 (purple) and then returned a different context to that used during habituation on Day 5-9 (burgundy). Horizontal lines indicate the median. There was a significant difference between testing days. (F, G) Freeze duration (F) and latency (G) in mice initially tested in a different context (Day 0, burgundy), then tested in the habituated (same) context on Day 1 (purple) and then returned to the original Day 0 context on Day 5-9 (burgundy). Horizontal lines indicate the median.

Consistent with the data shown in Figure 4, freezing was low when mice were tested in the same context as they were exposed to during habituation immediately following the habituation protocol (Day 0; 33.2 ± 7.9%; n=9 mice), but significantly increased when mice were tested in a different context the next day (Day 1; 79.8 ± 6.6%; n=9 mice). Importantly, freezing returned to low levels (29.6 ± 8.7%; n=9 mice) on Day 5–9 when mice were tested again in the same context as they were exposed to during habituation (Figure 5B, one-way ANOVA, F (2, 24)=12.89, p=0.0002). Despite this context-specific reversal in habituation, freeze duration was not significantly different between Day 0, 1 and 5–9 (Figure 5C, Kruskal-Wallis test, H=5.744, p=0.0566). Freeze latency also did not differ significantly across days (Figure 5D, Kruskal-Wallis test, H=2.231, p=0.3277).

We also tested the opposite situation, where mice were tested in a different context from the one used during habituation immediately following the habituation protocol. This led to high levels of freezing (Day 0; 87.1 ± 6.6%; n=9 mice), which was significantly reduced the following day when mice were tested in the same context as used during habituation (25.7 ± 4.9%; n=9 mice). Importantly, freezing returned to high levels (85.2 ± 8.1%) on Day 5-9 when mice were tested in the different context from that used during habituation (Figure 5E, one-way ANOVA, F(2, 24)=27.53, p < 0.0001). Freeze duration on Day 1 was significantly lower than on Day 0 in these mice (Figure 5F, Kruskal-Wallis test, H=9.275, p=0.0097; Dunn’s post hoc test, Day 0 vs. Day 1: p=0.0084), although freeze latency was unchanged across the different testing days (Figure 5G, Kruskal-Wallis test, H=0.3702, p=0.8308). Together, these results demonstrate that mice robustly associate habituation with the context in which it occurred, indicating that habituation is gated by specific contextual cues and is reversible.

## Discussion

In this study, we describe a behavioural protocol that rapidly induces habituation of freezing responses to slowly sweeping visual stimuli in mice. Consistent with recent work using looming stimuli, we show that it is possible to rapidly habituate mice to slowly sweeping visual stimuli within minutes^15,18^. Using this paradigm, we found that environmental context plays a critical role in determining the impact of habituation on visually evoked defensive behaviours. Specifically, we found that changing the context led to a significant and reversible reduction in habituation of freezing responses to slowly sweeping stimuli simulating an aerial predator cruising overhead. These data indicate that non-associative forms of learning, such as habituation of innate defensive behaviours, also depend on context. In doing so, our findings provide new insights into how context shapes habituation of an evolutionarily conserved defensive system.

We induced robust habituation to sweeping visual stimuli within minutes. Our habituation protocol was based on a recent study^18^, that achieved rapid suppression of escape responses to a looming visual stimulus by gradually increasing the stimulus-background contrast across repeated presentations. This work highlights how changes in stimulus salience can facilitate plasticity within innate defensive circuits. We adapted this contrast ramping approach to sweeping stimuli, which elicit freezing rather than escape, and observed a comparable reduction in defensive behaviour. This cross-stimulus generality suggests that gradual contrast enhancement may be a broadly effective strategy for attenuating innate defensive responses to visually threatening stimuli. One possibility is that the progressive increase in visual salience allows time for perceptual re-evaluation and thus diminishes the perceived threat level, allowing animals to reclassify the stimulus as non-threatening. Alternatively, contrast ramping may engage neural circuits involved in sensory prediction or top-down inhibition, engaging circuits that prevent overactivation of midbrain defence pathways^29,30^. Together, these findings demonstrate that dynamic manipulation of sensory input can serve as a powerful tool for probing and shaping innate behavioural responses.

Previous studies on innate fear responses suggest that these responses are tightly conserved, species-specific and evolutionarily tuned to specific ecological threats^8,30–33^. According to this view, the innate defensive behaviour elicited depends on the stimulus conditions, with different behaviours generated by overhead movement, looming expansion, or predator-like silhouettes, each presumably generated by hard-wired neural circuits designed to ensure survival. Consistent with this idea, earlier work has shown that habituation of innate defensive responses depends on critical stimulus parameters, such as the surface area and shape of the threatening visual stimulus, the contrast between the stimulus and background and stimulus speed, where increasing speed can reinstate defensive behaviour even after habituation^13,19^. Based on these observations, one might expect that changing the direction of a sweeping stimulus would also disrupt habituation. In contrast to this view, our results reveal that habituation to sweeping visual stimuli is not dependent on stimulus direction. Specifically, mice that were habituated to sweeping stimuli in one direction displayed comparable reductions in freezing when tested with stimuli moving in the orthogonal direction. One explanation for this is that certain global stimulus features, such as overall trajectory or direction, may be less critical in defining threat salience of sweeping stimuli, with the brain encoding classes of threat stimuli at a broader representational level capable of generalising across changes in direction of the same stimulus type. Alternatively, as mice freely move in the arena during the habituation session, they will be exposed to the habituating stimulus from multiple directions. This may diminish the salience of direction as a distinguishing factor, explaining why changing the stimulus direction does not restore defensive behaviours.

Theories of habituation, such as Wagner’s model of stimulus processing, provide a framework for understanding this context dependency^34^. In this model, repeated stimulus exposure activates a representation in short-term memory (STM), inhibiting further responses through a priming mechanism. Over time, associative links may form between the stimulus and contextual cues, which could be stored in long-term memory (LTM). When the stimulus is later encountered in the same context, these contextual cues reactivate the stimulus representation in STM, sustaining the habituated response. In a novel context, where such associations are absent, this retrieval may fail and the animal may revert to the initial defensive behaviour. Similarly, comparator models propose that the brain forms internal models of stimuli based on prior experience, which may incorporate contextual elements. If incoming sensory inputs (including context) match the internal model, behavioural responses are suppressed, whereas mismatches lead to the recurrence of defensive responses^35^. These theoretical models align with our observation that changing the context after habituation reinstated freezing, while testing in the same context preserved the habituated state.

While the superior colliculus (SC) is known to be involved in the rapid detection of threatening visual stimuli and plays a central role in initiating innate defensive responses such as freezing and escape^36^, the context dependency of habituation likely involves higher-order brain regions that mediate contextual learning and emotional regulation. In particular, the hippocampus and the amygdala have been implicated in encoding contextual associations and orchestrating defensive behaviours^37–39^. The amygdala is known as a key structure for initiating freezing responses and receives threat-related visual input via excitatory projections from the SC through the lateral posterior thalamic nucleus^29^. The hippocampus, by contrast, encodes contextual information and exerts top-down control of amygdala activity through direct projections^37–39^. Previous work has shown that the strength of the hippocampal projection to the basolateral amygdala, particularly from the ventral CA1, increases during fear learning, enabling context-specific retrieval of aversive memories^37,40^. In addition, reactivation of hippocampal engram cells active in a context that was previously associated with threat can drive the activation of amygdala engram neurons, allowing the animal to express fear ^39^. Although habituation involves a reduction rather than an enhancement of fear responses, these findings suggest that hippocampal–amygdala circuits are crucial for linking context to threat relevance. This may suggest that habituation downregulates hippocampal–amygdala circuitry driven by a specific context, suppressing fear expression.

In humans, contextual modulation plays a critical role in shaping fear responses, and its disruption is a hallmark of anxiety-related disorders. In post-traumatic stress disorder (PTSD), for example, environmental cues associated with prior trauma can evoke intense defensive reactions even in the absence of a direct threat^41^. These maladaptive responses are thought to arise from impaired contextual discrimination, where fear is not appropriately suppressed in safe contexts^42^. Neuroimaging studies have consistently shown hyperactivity of the amygdala and altered hippocampal function in individuals with PTSD, reflecting a breakdown in the neural circuits involved in contextual fear regulation^43–45^. While habituation in humans is less well understood, studies show that defensive responses can reappear in novel or trauma-associated contexts, suggesting that, as in rodents, contextual cues play a critical role in shaping behaviour^46,47^. Consistent with this, our findings demonstrate that environmental context influences the expression of defensive behaviours beyond the effects of stimulus repetition. Together, these observations highlight context as a key factor in modulating both innate and learned fear responses.

In conclusion, our findings establish a rapid and robust paradigm for habituating innate defensive behaviours in mice, while demonstrating that habituation is modulated by contextual factors. This highlights a surprising degree of flexibility in what is traditionally considered a hard-wired and fixed defensive system. Our study also presents a tractable model for examining how threat responses are dynamically influenced by experience. Given that deficits in habituation have been linked to neuropsychiatric conditions such as PTSD, autism spectrum disorder, and Attention-Deficit / Hyperactivity Disorder (ADHD)^48,49^, understanding how the brain flexibly suppresses defensive behaviours in safe environments may offer insights into the neural basis of maladaptive fear and hypervigilance in these disorders. Future studies investigating habituation and its context dependence at the cellular and circuit level may help uncover how the brain balances innate threat detection with learned safety - a fundamental process for both animal survival and human mental health.

## Materials and Methods

### Experimental Animals

A total of 26 mice were used in this study (age 6-8 weeks at the start of the experiment). All mice were housed in a 12:12 reversed light/dark cycle and were acclimatised for one week to adjust to this light cycle before the commencement of experiments. Two days prior to the experiments, mice were separated into individual cages with enrichment (enrichment roll or wood block). Food and water were provided ad libitum throughout the study.

### Experimental Setup

The behavioural arena used in this study was a rectangular opaque box with dimensions of 47 cm in length, 36 cm in width and 30 cm in height. A Samsung P2050 LCD monitor (38 cm x 27 cm, 300 cd/m2 brightness) was used to present visual stimuli. The monitor was placed above the arena, parallel to the floor. An LED light, mounted on the arena wall, was used to indicate the start and end of visual stimulus presentation, providing a reference point for video analysis. Mice could not see this LED light. Mouse behaviour was recorded using a PlayStation 3 Eye Camera at 75 frames per second in MP4 format. The camera was placed between the LCD monitor and the arena wall, capturing both the LED light and behavioural responses. In addition to a plain, undecorated arena, in some experiments the arena was decorated with two distinct contexts, which differed visually and tactually: Context A featured four laminated visual cues (horizontal lines, hollow square dots, vertical lines and crossed lines) attached to the arena walls using Blu-Tack. Context B had bare walls, but the floor was covered with bubble wrap securely taped to the arena surface.

### Stimulus properties

The sweeping stimulus used during experiments consisted of a 2.5 cm diameter black disk which traversed the monitor over 4 seconds from one corner to the opposite corner, moving at a velocity of 11.7 cm/s (Figure 1A). Stimuli were generated using Psychtoolbox v3 (Brainard, 1997) in conjunction with MATLAB R2019 software (MathWorks, Inc., Natick, MA).

### Behavioural Experiments

Mice in the control group were placed in the arena with the LCD monitor set to a white background and allowed to explore the arena for 9 minutes. This ensures that control mice had the same amount of time to explore the test arena as the habituation group, minimising any differences due to familiarity with the arena. After this exploration period, control mice were briefly transferred back to their home cage for approximately 2 minutes to avoid exposure to screen flashes during initialisation of the Psychtoolbox test sweep program. When the test sweep program was ready, mice were returned to the arena for testing. The full-contrast test sweeping stimulus was manually initiated when mice entered the centre of the arena, marked by four dots on the floor (Figure 1A). During each testing session, mice were presented with up to three sweeping stimuli separated by at least 90 seconds. Not all mice entered the centre of the arena three times within the 30-minute duration of the testing period. Hence, the total number of trials/mice per group differs between experiments. Mice were only tested once per day, except when a mouse did not enter the arena centre within 20 minutes of the onset of the testing phase. In this case, the experiment was terminated and the mouse was retested later that day.

Mice exposed to the habituation protocol were placed in the arena with the LCD monitor initially displaying a dark background. Mice were then exposed to 130 sweeping black disks, which traversed the monitor screen diagonally in 4 seconds. The protocol began with sweeping black disks presented against a dark background (grey level 2 in MATLAB), with the background incrementally increasing in lightness by two grey levels after each stimulus until the highest contrast condition was reached (grey level 255 in MATLAB), after which the sweeping black disk stimulus was presented against a white background three times at the highest contrast level. Given that each sweep lasted 4 seconds, the habituation protocol lasted approximately 9 minutes. After the habituation protocol, mice were briefly transferred back to their home cage for approximately 2 minutes during initialisation of the Psychtoolbox test sweep program, then returned to the arena for testing using the same full-contrast test sweeping stimuli (as explained above).

### Manual and Computational Analysis of Behavioural Data

Experiment videos were manually reviewed to identify and classify behavioural responses to the sweeping stimulus. Adobe Premiere Pro 2024 was used to manually determine the onset and offset of freezing behaviour by frame-by-frame inspection of the video. Freezing was classified as a sudden reduction in speed, where mouse movement decreased to 2 cm/s or less for a minimum of 0.5 seconds. Freeze offset was defined as the frame where the mouse resumed movement. During this period, mice must not exhibit any voluntary movements such as grooming, rearing or sniffing. The absence of these behaviours was crucial for a behaviour to be classified as freezing. The analysis window for freezing was constrained to the duration of the stimulus, which was 4 seconds.

To determine freeze duration, the frame number at which a mouse initially froze was subtracted from the frame number at which the mouse resumed movement. To determine freeze latency, the frame number at the onset of the sweeping stimulus (as detected by LED onset) was subtracted from the frame number at which the mouse initially froze. The difference in frame number was then converted to time in seconds to determine freeze duration and latency based on the frame rate (75 frames per second). DeepLabCut and MATLAB were used in conjunction with manual analysis. Automated analysis using DeepLabCut was primarily used to double-check responses when manual analysis was ambiguous and to plot raster and trajectory figures.

### Statistical tests

All statistical analysis was performed using GraphPad Prism version 10.3.1 (GraphPad Software, Inc). For the calculation of the average percentage of freezing, the average freezing response was calculated for each mouse across all trials and treated as a continuous variable. Data normality was assessed using both the Shapiro–Wilk test and the D’Agostino–Pearson omnibus test. For comparisons between two groups, unpaired t-tests were used if data were normally distributed; otherwise, the nonparametric Mann–Whitney test was applied. For comparisons involving more than two groups, one-way ANOVA was used for normally distributed data, followed by Tukey’s multiple comparisons test when significant differences were found. If the data were not normally distributed, the Kruskal–Wallis test was used, followed by Dunn’s multiple comparisons test for post-hoc analysis when appropriate. For all the analyses, statistical significance was defined as p < 0.05. Statistical comparisons are indicated in the figures by asterisks: p < 0.05 (*), p < 0.01 (**), p < 0.001 (***), and p < 0.0001 (****). Normally distributed data are reported as mean ± standard error of the mean (SEM), while non-normally distributed data are reported as median with interquartile range (IQR).

## Acknowledgments

This work was supported by the Australian Research Council (CE140100007), the Australian National University and Monash University. We thank Ehsan Arabzadeh for providing mice used in this study.

## Author Contributions

Conceptualisation and methodology: S.G. and G.J.S.; Investigation and analysis: H.Q. and A.T.; Writing: H.Q., S.G. and G.J.S.; Supervision: S.G. and G.J.S.

## Data availability

The data that support the findings of this study are available from the corresponding authors upon reasonable request.

## References

1. Tseng, Y.-T., Schaefke, B., Wei, P., and Wang, L. (2023). Defensive responses: behaviour, the brain and the body. Nature Reviews Neuroscience 24, 655–671. 10.1038/s41583-023-00736-3.

2. Campos-Cardoso, R., and Cummings, K.A. (2025). Threat learning: Avoiding danger in the first place. Current Biology 35, R177–R180. 10.1016/j.cub.2025.01.039.

3. Blanchard, D., and Blanchard, R. (2008). Chapter 2.4 Defensive behaviors, fear, and anxiety. Handbook of Behavioral Neuroscience 17, 63–79. 10.1016/S1569-7339(07)00005-7.

4. Kawai, N., Kono, R., and Sugimoto, S. (2004). Avoidance learning in the crayfish (Procambarus clarkii) depends on the predatory imminence of the unconditioned stimulus: a behavior systems approach to learning in invertebrates. Behavioural Brain Research 150, 229–237. 10.1016/S0166-4328(03)00261-4.

5. Blanchard, R.J., and Blanchard, D.C. (1989). Attack and defense in rodents as ethoexperimental models for the study of emotion. Prog Neuropsychopharmacol Biol Psychiatry 13 *Suppl*, S3-14. 10.1016/0278-5846(89)90105-x.

6. Wu, Q., and Zhang, Y. (2023). Neural Circuit Mechanisms Involved in Animals’ Detection of and Response to Visual Threats. Neuroscience Bulletin 39, 994–1008. 10.1007/s12264-023-01021-0.

7. Fanselow, M.S., and Lester, L.S. (2013). A functional behavioristic approach to aversively motivated behavior:: Predatory imminence as a determinant of the topography of defensive behavior. In Evolution and learning, (Psychology Press), pp. 185–212.

8. Mancienne, T., Marquez-Legorreta, E., Wilde, M., Piber, M., Favre-Bulle, I., Vanwalleghem, G., and Scott, E.K. (2021). Contributions of Luminance and Motion to Visual Escape and Habituation in Larval Zebrafish. Front Neural Circuits 15, 748535. 10.3389/fncir.2021.748535.

9. Silva, B.A., Gross, C.T., and Gräff, J. (2016). The neural circuits of innate fear: detection, integration, action, and memorization. Learn Mem 23, 544–555. 10.1101/lm.042812.116.

10. De Franceschi, G., Vivattanasarn, T., Saleem, A.B., and Solomon, S.G.(2016). Vision Guides Selection of Freeze or Flight Defense Strategies in Mice. Curr Biol 26, 2150–2154. 10.1016/j.cub.2016.06.006.

11. Solomon, S.G., Janbon, H., Bimson, A., and Wheatcroft, T. (2023). Visual spatial location influences selection of instinctive behaviours in mouse. Royal Society Open Science 10, 230034. 10.1098/rsos.230034.

12. de Vries, Saskia E.J., and Clandinin, Thomas R. (2012). Loom-Sensitive Neurons Link Computation to Action in the *Drosophila* Visual System. Current Biology 22, 353–362. 10.1016/j.cub.2012.01.007.

13. Yilmaz, M., and Meister, M. (2013). Rapid Innate Defensive Responses of Mice to Looming Visual Stimuli. Current Biology 23, 2011–2015. 10.1016/j.cub.2013.08.015.

14. Haimson, B., and Mizrahi, A. (2025). Integrating innate and learned behavior through brain circuits. Trends in Neurosciences 48, 319–329. 10.1016/j.tins.2025.03.002.

15. Mederos, S., Blakely, P., Vissers, N., Clopath, C., and Hofer, S.B. (2025). Overwriting an instinct: Visual cortex instructs learning to suppress fear responses. Science 387, 682–688. doi:10.1126/science.adr2247.

16. Narushima, M., Agetsuma, M., and Nabekura, J. (2022). Development and experience-dependent modulation of the defensive behaviors of mice to visual threats. The Journal of Physiological Sciences 72, 5. 10.1186/s12576-022-00831-7.

17. Carroll, J.N., Myers, B., and Vaaga, C.E. (2025). Repeated presentation of visual threats drives innate fear habituation and is modulated by threat history and acute stress exposure. Stress 28, 2489942. 10.1080/10253890.2025.2489942.

18. Lenzi, S.C., Cossell, L., Grainger, B., Olesen, S.F., Branco, T., and Margrie, T.W. (2022). Threat history controls flexible escape behavior in mice. Current Biology 32, 2972–2979.e2973. 10.1016/j.cub.2022.05.022.

19. Tafreshiha, A., van der Burg, S.A., Smits, K., Blömer, L.A., and Heimel, J.A. (2021). Visual stimulus-specific habituation of innate defensive behaviour in mice. J Exp Biol 224. 10.1242/jeb.230433.

20. Thompson, R.F., and Spencer, W.A. (1966). Habituation: A model phenomenon for the study of neuronal substrates of behavior. Psychological Review 73, 16–43. 10.1037/h0022681.

21. Liu, X., Lai, J., Han, C., Zhong, H., Huang, K., Liu, Y., Zhu, X., Wei, P., Tan, L., Xu, F., and Wang, L. (2025). Neural circuit underlying individual differences in visual escape habituation. Neuron. 10.1016/j.neuron.2025.04.018.

22. Byrne, J.H. (2013). Chapter 47 - Learning and Memory: Basic Mechanisms. In Fundamental Neuroscience (Fourth Edition), L.R. Squire, D. Berg, F.E. Bloom, S. du Lac, Ghosh, and N.C. Spitzer, eds. (Academic Press), pp. 1009–1027. 10.1016/B978-0-12-385870-2.00047-0.

23. Maren, S., Phan, K.L., and Liberzon, I. (2013). The contextual brain: implications for fear conditioning, extinction and psychopathology. Nature Reviews Neuroscience 14, 417–428. 10.1038/nrn3492.

24. Chu, A., Gordon, N.T., DuBois, A.M., Michel, C.B., Hanrahan, K.E., Williams, D.C., Anzellotti, S., and McDannald, M.A. (2024). A fear conditioned cue orchestrates a suite of behaviors in rats. Elife 13. 10.7554/eLife.82497.

25. Izquierdo, I., Furini, C.R.G., and Myskiw, J.C. (2016). Fear Memory. Physiological Reviews 96, 695–750. 10.1152/physrev.00018.2015.

26. Pavlov, P.I. (2010). Conditioned reflexes: An investigation of the physiological activity of the cerebral cortex. Ann Neurosci 17, 136–141. 10.5214/ans.0972-7531.1017309.

27. Hassien, A.M., Shue, F., Bernier, B.E., and Drew, M.R. (2020). A mouse model of stress-enhanced fear learning demonstrates extinction-sensitive and extinction-resistant effects of footshock stress. Behav Brain Res 379, 112391. 10.1016/j.bbr.2019.112391.

28. Broersen, R., Thompson, G., Thomas, F., and Stuart, G.J. (2025). Binocular processing facilitates escape behavior through multiple pathways to the superior colliculus. Current Biology 35, 1242–1257.e1249. 10.1016/j.cub.2025.01.066.

29. Wei, P., Liu, N., Zhang, Z., Liu, X., Tang, Y., He, X., Wu, B., Zhou, Z., Liu, Y., Li, J., et al. (2015). Processing of visually evoked innate fear by a non-canonical thalamic pathway. Nature Communications 6, 6756. 10.1038/ncomms7756.

30. Baier, F., Reinhard, K., Nuttin, B., Sans-Dublanc, A., Liu, C., Tong, V., Murmann, J.S., Wierda, K., Farrow, K., and Hoekstra, H.E. (2025). The neural basis of species-specific defensive behaviour in Peromyscus mice. Nature. 10.1038/s41586-025-09241-2.

31. Bolles, R.C. (1971). CHAPTER 3 - Species-Specific Defense Reactions. In Aversive Conditioning and Learning, F.R. Brush, ed. (Academic Press), pp. 183–233. 10.1016/B978-0-12-137950-6.50008-0.

32. Blanchard, R.J., and Blanchard, D.C. (1971). Defensive reactions in the albino rat. Learning and Motivation 2, 351–362. 10.1016/0023-9690(71)90016-6.

33. Dielenberg, R.A., and McGregor, I.S. (2001). Defensive behavior in rats towards predatory odors: a review. Neuroscience & Biobehavioral Reviews 25, 597–609. 10.1016/S0149-7634(01)00044-6.

34. Wagner, A.R. (2014). Habituation and memory. In Mechanisms of learning and motivation, (Psychology Press), pp. 53–82.

35. Sokolov, E.N. (1963). Higher nervous functions; the orienting reflex. Annu Rev Physiol 25, 545–580. 10.1146/annurev.ph.25.030163.002553.

36. Shang, C., Chen, Z., Liu, A., Li, Y., Zhang, J., Qu, B., Yan, F., Zhang, Y., Liu, W., Liu, Z., et al. (2018). Divergent midbrain circuits orchestrate escape and freezing responses to looming stimuli in mice. Nature Communications 9, 1232. 10.1038/s41467-018-03580-7.

37. Kim, W.B., and Cho, J.-H. (2020). Encoding of contextual fear memory in hippocampal– amygdala circuit. Nature Communications 11, 1382. 10.1038/s41467-020-15121-2.

38. Terranova, J.I., Yokose, J., Osanai, H., Marks, W.D., Yamamoto, J., Ogawa, S.K., and Kitamura, T. (2022). Hippocampal-amygdala memory circuits govern experience-dependent observational fear. Neuron 110, 1416–1431.e1413. 10.1016/j.neuron.2022.01.019.

39. Josselyn, S.A., Köhler, S., and Frankland, P.W. (2015). Finding the engram. Nat Rev Neurosci 16, 521–534. 10.1038/nrn4000.

40. Jimenez, J.C., Berry, J.E., Lim, S.C., Ong, S.K., Kheirbek, M.A., and Hen, R. (2020). Contextual fear memory retrieval by correlated ensembles of ventral CA1 neurons. Nature Communications 11, 3492. 10.1038/s41467-020-17270-w.

41. Liberzon, I., and Sripada, C.S. (2008). The functional neuroanatomy of PTSD: a critical review. Prog Brain Res 167, 151–169. 10.1016/s0079-6123(07)67011-3.

42. Grillon, C. (2002). Startle reactivity and anxiety disorders: aversive conditioning, context, and neurobiology. Biological Psychiatry 52, 958–975. 10.1016/S0006-3223(02)01665-7.

43. Shin, L.M., and Liberzon, I. (2010). The Neurocircuitry of Fear, Stress, and Anxiety Disorders. Neuropsychopharmacology 35, 169–191. 10.1038/npp.2009.83.

44. Liberzon, I., Taylor, S.F., Amdur, R., Jung, T.D., Chamberlain, K.R., Minoshima, S., Koeppe, R.A., and Fig, L.M. (1999). Brain activation in PTSD in response to trauma-related stimuli. Biol Psychiatry 45, 817–826. 10.1016/s0006-3223(98)00246-7.

45. Tsoory, M.M., Vouimba, R.M., Akirav, I., Kavushansky, A., Avital, A., and Richter-Levin, G. (2007). Amygdala modulation of memory-related processes in the hippocampus: potential relevance to PTSD. In Progress in Brain Research, E.R. De Kloet, M.S. Oitzl, and E. Vermetten, eds. (Elsevier), pp. 35–51. 10.1016/S00796123(07)67003-4.

46. LaBar, K.S., and Phelps, E.A. (2005). Reinstatement of conditioned fear in humans is context dependent and impaired in amnesia. Behav Neurosci 119, 677–686. 10.1037/0735-7044.119.3.677.

47. Andreatta, M., and Pauli, P. (2021). Contextual modulation of conditioned responses in humans: A review on virtual reality studies. Clinical Psychology Review 90, 102095. 10.1016/j.cpr.2021.102095.

48. McDiarmid, T.A., Bernardos, A.C., and Rankin, C.H. (2017). Habituation is altered in neuropsychiatric disorders—A comprehensive review with recommendations for experimental design and analysis. Neuroscience & Biobehavioral Reviews 80, 286–305. 10.1016/j.neubiorev.2017.05.028.

49. Jamal, W., Cardinaux, A., Haskins, A.J., Kjelgaard, M., and Sinha, P. (2021). Reduced Sensory Habituation in Autism and Its Correlation with Behavioral Measures. Journal of Autism and Developmental Disorders 51, 3153–3164. 10.1007/s10803-020-04780-1.

